# A novel virulence phenotype rapidly assesses *Candida* fungal pathogenesis in healthy and immunocompromised *Caenorhabditis elegans* hosts

**DOI:** 10.1101/370403

**Authors:** Dorian J. Feistel, Rema Elmostafa, Nancy Nguyen, McKenna Penley, Levi Morran, Meleah A. Hickman

## Abstract

The yeast *Candida albicans* is an opportunistic pathogen of humans, meaning that despite commensal interactions with its host, it can transition to a harmful pathogen. While *C. albicans* is the predominant species isolated in the human mycobiome and implicated in fungal infection, infections due to non-albicans *Candida* species are rapidly rising. Studying the factors that contribute to virulence is often challenging and frequently depends on many contexts including host immune status and pathogen genetic background. Here, we utilize the nematode *Caenorhabditis elegans* as a perspicuous and efficient model host system to study fungal infections of *Candida* pathogens. We find that in addition to reducing lifetime host survival, exposure to *C. albicans* results in delayed reproduction, which significantly reduced lineage growth over multiple generations. Furthermore, we assessed fungal pathogen virulence in *C. elegans* hosts compromised for innate immune function and detected increased early mortality, reduced brood sizes and delayed reproduction relative to infected healthy hosts. Importantly, by assessing virulence in both healthy and immunocompromised host backgrounds we reveal the pathogen potential in non-*albicans Candida* species. Taken together, we present a novel lineage growth assay to measure reduction in host fitness associated with fungal infection and demonstrate significant interactions between pathogen and host immune function that contribute to virulence.

## Importance

Opportunistic pathogens are commensals capable of causing disease and are serious threats to human health. It is critical to understand the mechanisms and host contexts under which opportunistic pathogens become virulent. In this work, we present a novel assay to quickly and quantitatively measure pathogen virulence in healthy and immunocompromised hosts. We found that *Candida* species, the most prominent fungal opportunistic pathogens of humans, decrease host fitness by reducing survival and impacting host reproduction. Most importantly, by measuring virulence in hosts that have intact or compromised immune function, we can reveal the pathogenic potential of opportunistic fungal pathogens.

## Introduction

*Candida* species are commensals of the human gastrointestinal microbiota and various other niches in the human body [1]. Despite their commensal existence in many humans, *Candida* species also account for the most fungal infections in the world and are the fourth most prevalent cause of all nosocomial blood stream infections [2,3]. The severity of fungal infection is often dependent on the immune status of the host, with superficial infections occurring in healthy individuals and bloodstream infections in immunocompromised hosts [4-8]. The majority of these infections are associated with *Candida albicans*, however other non-*albicans Candida* species are increasingly becoming implicated in fungal infections [9,10].

Several host models have been employed to better understand fungal infection and disease progression. These experimental systems include mouse, zebrafish, nematode, wax moth and human *ex vivo* models have been specifically developed to study *Candida* infections [11]. Most often, virulence phenotypes are limited to survival outcomes or tissue-specific fungal burden [12], the latter of which requires animal sacrifice and cannot address the long-term consequence of fungal disease. Furthermore, animal mortality and sacrifice, particularly in vertebrate models, constrain the number of infected individuals to study and thus reduce the quantitative robustness of these experiments. Nonetheless, these studies provide valuable insight into *Candida* infection and disease and have uncovered important roles for innate immune function of hosts [13-15], specific *C. albicans* genes regulating virulence [16,17] and differences between genetic backgrounds within and between *Candida* species [18].

Other major limitations for most studies of *Candida* infection virulence include the time-intensive nature of tracking virulence phenotypes in vertebrate systems such as mice, the genetic intractability of zebrafish and wax moth larvae and the context-independent nature of *ex vivo* models [4]. As such, most studies focus specifically on *C. albicans* virulence while many non-albicans *Candida* species are evaluated to a lesser degree. The nematode *Caenorhabditis elegans* has been developed as a model for *Candida* infection to overcome these limitations [19,20] and to screen compounds for antifungal activity [21]. *C. elegans* has proven useful for studying host-microbe interactions [22] because many fungal and bacterial pathogens that cause illness in humans also cause disease in *C. elegans* [23]. Infecting *C. elegans* is relatively simple and often done by replacing or incorporating its standard laboratory food *E. coli* with a desired pathogen which colonizes the gut and causes disease [23]. Furthermore, the innate immune system in *C. elegans* includes the *SEK-1* gene which encodes a MAP Kinase Kinase (MAPKK) that is homologous to the MKK3/6 and MKK4 family of mammalian MAPKKs and activates the *C. elegans* p38 MAP kinase ortholog [24]. This pathway has been suggested to be an ancient and conserved component of *C. elegans’* immune response to pathogens [25]. As such, the *sek-1* mutation increases susceptibility to microbial colonization [26], including infection with many *Candida* species [27] and is necessary to induce the appropriate antifungal immune defense [28].

In addition to utilizing innate immune system mutants, most studies measure host mortality in temperature sensitive sterile mutants [17,21,29,30] to more easily track mortality in founder populations since *C. elegans* has a rapid lifecycle and large brood sizes [31,32]. The effects on host reproduction are often overlooked in favor of examination of mortality rates when assessing a pathogen’s virulence [33]. Yet, it is important to remember that virulence can be broadly measured as any reduction in host fitness resulting from interactions between a pathogen and its host [34-36]. In this work, we assessed virulence of *Candida* species in *C. elegans* hosts using a novel measure of host fitness that incorporated both host survival and fecundity. We found that fungal pathogens reduced host fitness by delaying reproduction, resulting in long-term consequences for population growth in both healthy and immunocompromised host backgrounds. Using this novel measure, we characterized virulence phenotypes for three non-*albicans Candida* species; *C. dubliniensis, C. tropicalis*, and *C. parapsilosis* in both healthy and immunocompromised *C. elegans* hosts. Our studies demonstrate that differences in virulence can be identified between pathogen species and that pathogenic potential is often revealed when host immune function is diminished.

## Methods

### Strains and media

For this study, the fungal pathogens *C. albicans* (SC5314 [37]), *C. dubliniensis* (Wu284 [38]), *C. tropicalis* (ATCC 22109), and *C. parapsilosis* (ATCC 22109) strains were used. *C. elegans* N2 Bristol (WT) and a *sek-1* mutant derivative [24] were used to test host survival, fecundity and population growth. *C. elegans* populations were maintained at 20°C on 100mm petri dishes with 25 mL of lite nematode growth medium (NGM, US Biological) with *E. coli* OP50 as a food source. Nematodes were transferred to a newly seeded *E. coli* plate every 3-4 days. For survival, fecundity and population growth assays, NGM was supplemented with 0.08 g/L uridine, 0.08 g/L histidine, and 0.04 g/L adenine to facilitate growth of auxotrophic *C. albicans* strains and 0.2 g/L streptomycin sulfate to inhibit *E. coli* overgrowth so fungal strains could proliferate.

### Seeding NGM plates for survival, fecundity, and population growth assays

*Candida* strains and *E. coli* OP50 strains were inoculated in 3 mL of YPD or 5 mL of LB, respectively, and cultured at 30°C for 1-2 days. *Candida* culture densities were measured with a spectrophotometer and diluted to a final volume of 3.0 OD_600_ per mL (∼6 x 10^7^ cells per ml). *E. coli* cultures were pelleted and washed twice with 1 mL of ddH_2_O. The supernatant was removed and the pellet was centrifuged for 60 sec at maximum to remove any excess liquid. The pellet was weighed and suspended with sterilized water to a final volume of 200 mg/mL. A mix of 6.25 μL *E. coli*, 1.25 μL *Candida* was brought to a final volume of 50 μL with ddH_2_O. The entire 50 μL was spotted onto the center of a 35mm supplemented-NGM Lite agar plate and incubated at room temperature overnight before addition of eggs or transferring nematode. *E. coli* OP50 was used as a control at the same concentration described above.

### Egg preparation and synchronization for survival, fecundity and population growth assays

For survival, fecundity and population growth assays, approximately 100 nematodes at the L3/L4 stage were transferred to a 100mm NGM plate seeded with *E. coli* OP50 and maintained at 20°C for 2-3 days prior to the start of an experiment. On the first day of an experiment, these NGM plates were washed with M9 buffer and contents (live nematodes and eggs) transferred to 15 mL conical tube and pelleted by centrifugation (2 min at 1200 rpm). The pellet was re-suspended in a 1:4 bleach (5.25%) solution and transferred to a micro-centrifuge tube. The suspension was mixed via inversion for 90-120 sec and subsequently centrifuged (30 sec at 1500 rpm). The pellet was washed with 1 mL M9 buffer three consecutive times to remove excess bleach solution and brought to a final suspension with 500 μL M9 buffer. To determine the concentration of eggs, 10 μL was placed on a concaved slide and eggs counted and the egg suspension was diluted with M9 to a final concentration of 10 eggs/μL. All assays were treated equally on the first day (Day 0) by adding roughly 100 eggs to a treatment or *E. coli* plate (described above).

### Survival assays

This experimental procedure is a modified version of [39]. Briefly, 72 h after adding ∼50-75 nematode eggs to a plate, 40 adult nematodes were randomly selected and transferred to newly seeded plates with the same concentration of food as described above and incubated at 20°C. Every other day nematodes were transferred to freshly seeded plates until all nematode replicate populations went extinct. The number of living, dead, and censored worms were scored daily. Each survival assay was performed at least three independent times.

### Fecundity assays

For both fecundity and lineage expansion assays, 48 h after adding nematode eggs to a plate, a single L4 reproductively immature hermaphroditic nematode was randomly selected and transferred (6-10 independent biological replicates per treatment per block) to a newly seeded 35 mm petri plate containing either a treatment of *C. albicans* and *E. coli* or *E. coli* alone (described above) and incubated at 20°C. Nematodes were transferred to freshly seeded plates every 24 h for seven consecutive days. Eggs remained undisturbed on the plate and were incubated at 20°C for an additional 24 h to provide enough time to hatch, at which point the number of viable progeny per day was scored. Nematodes that died during the assay were scored as dead at the time of transfer. Nematodes that crawled off the plate or were otherwise unaccountable for were considered censored and excluded from the analysis.

### Lineage Expansion Assay

A single L4 reproductively immature hermaphroditic nematode was randomly selected and transferred (6 biological replicates per treatment) to a freshly seeded 100 mm treatment or *E. coli* plate (described above) containing a 6-fold increase in food. Five days later, each plate was washed with M9 buffer until the majority of nematodes were displaced and subsequently transferred to 15mL conical tubes. Tubes were placed at 4°C for 1h to allow the nematodes to settle at the bottom. All tubes were concentrated to a final volume of 10mL. Six, 20 μL samples were taken from each population and counted.

### Statistical analyses

All statistical analysis were performed with GraphPad Prism. Data sets were tested for normality using the D’Agostino & Pearson omnibus normality test. For comparisons across groups, one-way ANOVAs and post-hoc Tukey multiple comparison tests or Kruskal-Wallis and Dunn’s multiple comparison tests were performed.

## Results

### Exposure to C. albicans reduces multiple measures of C. elegans fitness

Pathogens affect host fitness by decreasing lifespan and/or reducing fecundity. To determine how *C. albicans* impacts *C. elegans* host fitness, we first measured nematode lifespan in the presence or absence of *C. albicans*. We observed a significant reduction in survival when *C. elegans* was reared on *C. albicans* compared to when it was reared in its absence (Figure 1, p < 0.0001 Log-rank test), consistent with previously published results [39]. 50% mortality was reached in eight days in nematode populations exposed to *C. albicans* compared to 16 days in unexposed nematode populations, indicating that *C. elegans* lifespan is substantially decreased by exposure to *C. albicans*.

**Figure 1:**
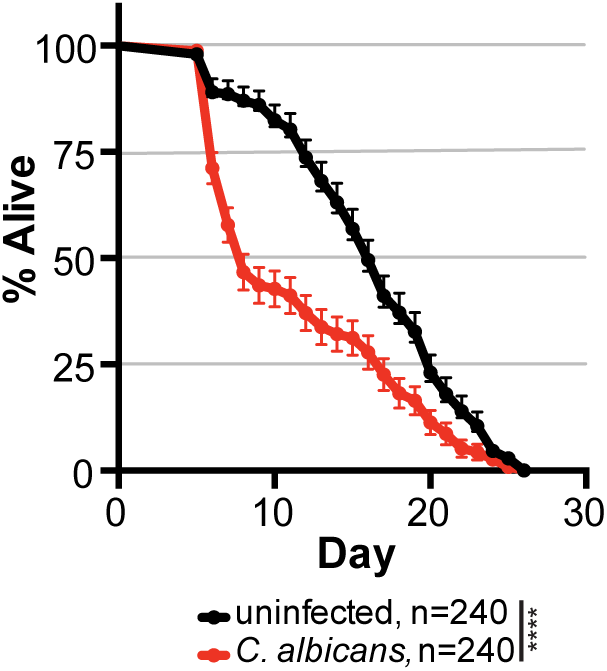
Reduced nematode survival associated with *C. albicans* infection. Survival curves for *C. elegans* populations that are either uninfected (exposed to just an *E. coli* food source, black) or when infected with *C. albicans* SC5314 (red). Mean data values are plotted with SEM error bars, number of worms analyzed (n) for each treatment is indicated in the legend. **** indicates p < 0.0001, Log-rank (Mantel-Cox) test.

While overall survival was reduced, nematode death was rarely observed earlier than 5 days post-infection. Importantly, adult nematodes produce the majority of their offspring before *C. albicans* infection reduces rates of survival. Given this information and that fecundity is a key component of fitness from an evolutionary perspective, we investigated whether *C. albicans* negatively impacts *C. elegans* reproduction. To test this, we counted the total number of viable progeny (i.e. brood size) produced in the first seven days of reproductive maturity from individual nematodes that were either unexposed or exposed to either heat-killed or live *C. albicans*. As shown in Figure 2A, exposure to *C. albicans* reduced the average number of viable progeny (253±7) compared to the unexposed control (285±4; *p* < 0.0002, Kruskal-Wallis test) by ∼11%. While statistically significant, the cost of *C. albicans* toward host brood size is modest. Furthermore, the brood size of nematodes exposed to heat-killed *C. albicans* (HK) was not detectably different from uninfected or live *C. albicans* treatments (Figure 2A).

**Figure 2.**
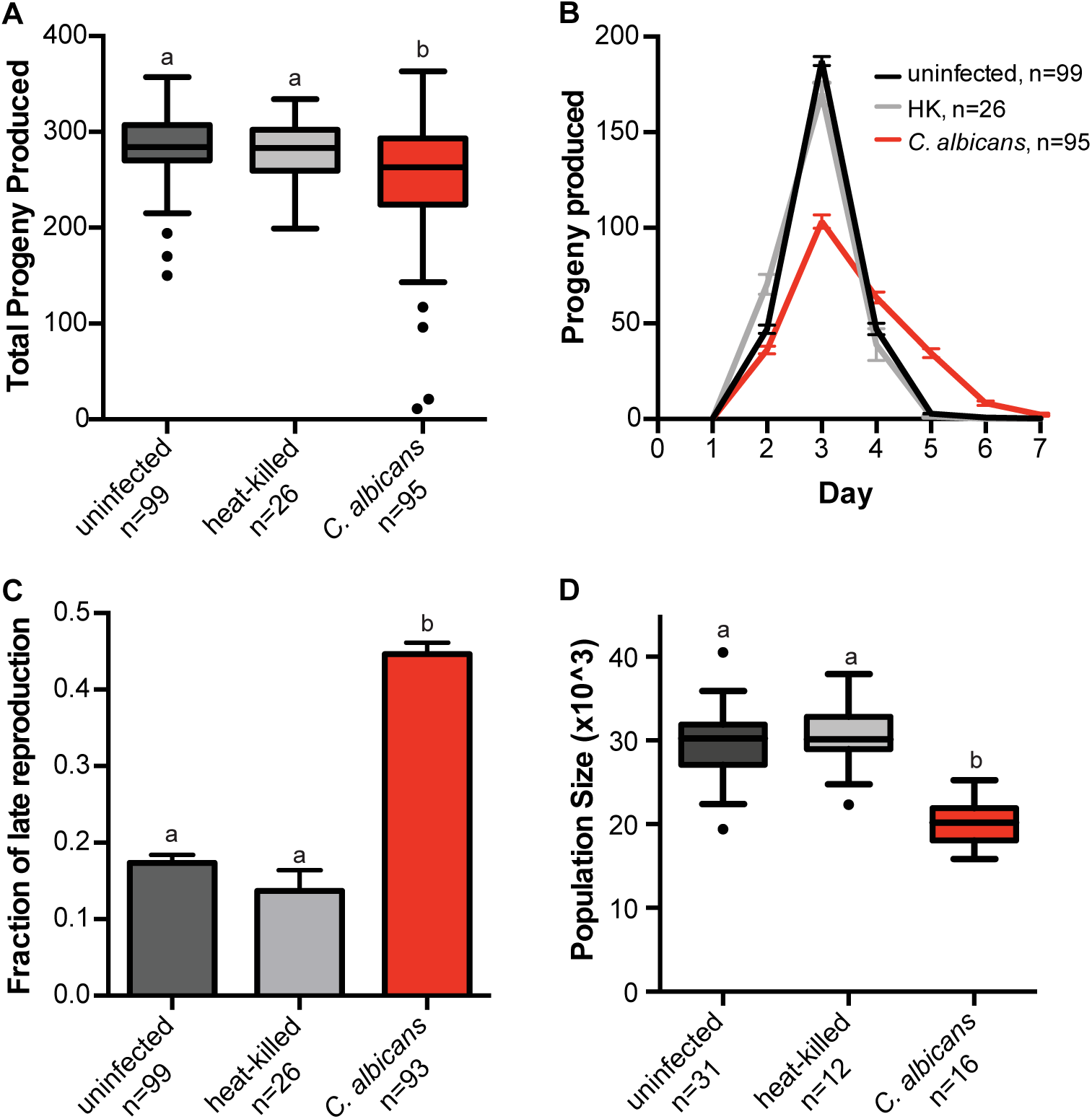
Live *C. albicans* reduces nematode fecundity. **A)** Box and whiskers plot of brood sizes for *C. elegans* (N2) exposed to *E. coli* food source alone (uninfected, black), heat-killed *C. albicans* (grey), or live *C. albicans* (red). Boxes indicate the 25-75th quar-tiles with median indicated. Error bars are the normalized range of the data and circles indicate outliers. **B)** Number of viable *C. elegans* progeny produced per day in uninfected, heat-killed, and live *C. albicans* treatments. Data represent the mean and SEM. **C)** The fraction of *C. elegans* progeny produced on Day4 or later during the ‘Late Reproductive’ window. Data represent the mean and SEM. **D)** Box and whiskers plot of average population size (representing the number of F1 and F2 progeny) produced from a single founder *elegans*. Boxes indicate the 25-75th quartiles with median indicated. Error bars are the normalized range of the data and circles indicate outliers. The number (n) of experimental samples analyzed is indicated for each treatment. Treatments that share letters are not significantly different, whereas treatments with differing letters are stastically significant, post-hoc Dunn’s multiple comparison test.

Intriguingly, we identified a significant delay in reproductive timing when we measured the number of progeny produced every 24 h during the host reproductive window. We observed a substantial reduction in reproduction on Day 3 in nematodes exposed to *C. albicans* (103±4) relative to unexposed hosts (187±3) or hosts associated with heat-killed *C. albicans* (Figure 2B; *p* < 0.0001 Kruskal-Wallis test) and increased progeny produced on Days 4-6 in nematodes exposed to *C. albicans* (Figure 2B). We calculated the total fraction of reproduction occurring in this ‘late’ window (Days 4-6) and identified a significant increase in the progeny produced late in *C. elegans* exposed to *C. albicans* compared to unexposed or heat-killed controls (Figure 2C; *p*=0.0001, Kruskal-Wallis test). Taken together, our data indicate that *C. albicans* severely delays and reduces reproduction in addition to impacting overall survival in *C. elegans*.

To investigate whether this reproductive delay observed in nematodes exposed to *C. albicans* had any long-term consequences, we took advantage that *C. elegans* are hermaphrodites and measured the total population produced from a single founder worm over multiple generations through lineage expansion assays. We find that the median population produced within 7 days from a single uninfected nematode is ∼30,250 F1 and F2 progeny (Figure 2D). Lineage expansion is significantly impacted in nematodes exposed to live *C. albicans*, with a median population size of ∼20,208 F1 and F2 progeny, a reduction of ∼33% compared to uninfected and heat-killed control treatments (Figure 2D, *p* < 0.0001 one-way ANOVA). We conclude that the delay in reproduction, coupled with the reduced brood size resulting from exposure to *C. albicans* severely impacts *C. elegans* evolutionary fitness.

### Immunocompromised hosts are susceptible to fungal infection

A leading host-related risk factor for invasive candidiasis is compromised immune function [40]. Given our novel results regarding fecundity in healthy *C. elegans* hosts, we were curious how exposure to *C. albicans* impacts nematode hosts with compromised immune function. To address this point, we utilized a *C. elegans* strain carrying a mutation in *SEK*-1, a well-conserved MAP kinase involved in the innate immune signaling cascade [24,26] and antifungal response [28]. Unlike healthy hosts, immunocompromised hosts show high rates of early (within 7 days) mortality in both uninfected and *C. albicans* treatments (Figure S1). *C. albicans* exposure reduced average brood size by nearly 50% (78±6) in immunocompromised hosts compared to uninfected (151±5) and heat-killed (127±13) controls (Figure 3A, p < 0.0001, Kruskal-Wallis test). It is important to note that in all treatments (unexposed, heat-killed, and live *C. albicans*), immunocompromised hosts produced significantly less progeny than healthy hosts (Table S1), but the largest reductions occur in immunocompromised hosts are exposed to live *C. albicans*.

**Figure 3:**
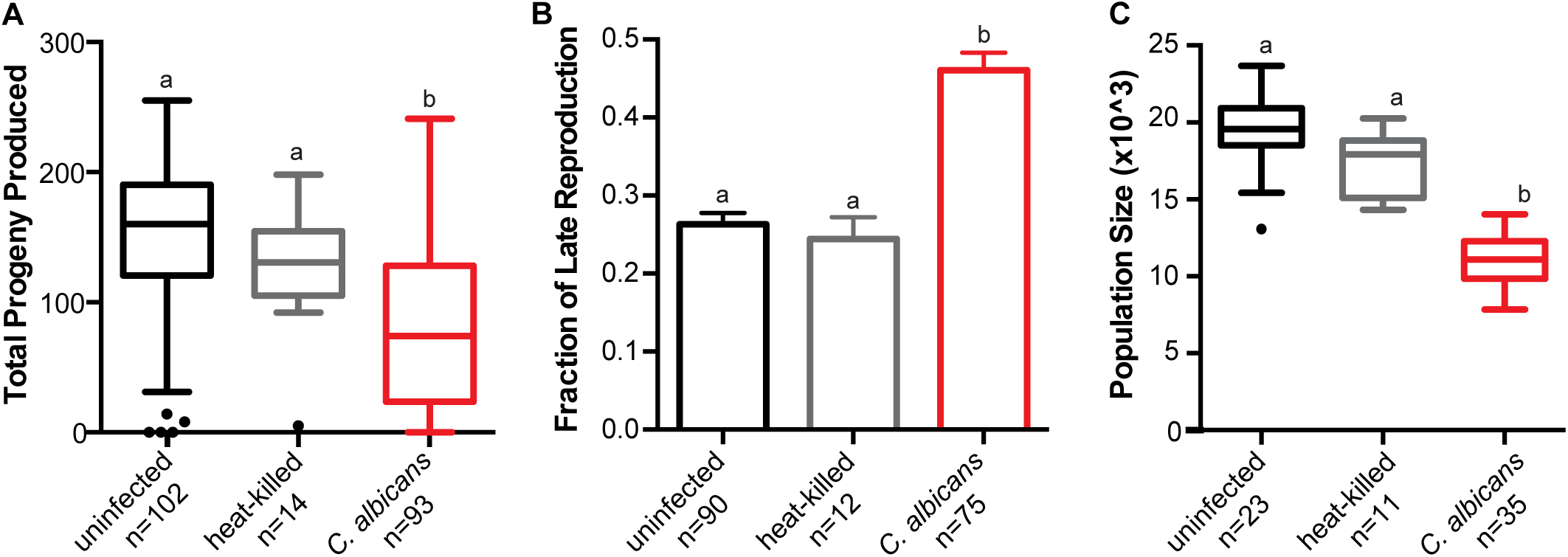
*C. albicans* severely impacts host fitness in immunocompromised nematodes. **A)** Box and whiskers plot of brood sizes for *C. elegans* (N2) exposure to *E. coli* food source alone (uninfected, black), heat-killed *C. albicans* (grey), or live *C. albicans* (red). Boxes indicate the 25-75th quartiles with median indicated. Error bars are the normalized range of the data and circles indicate outliers. **B)** The fraction of *C. elegans* progeny produced on Day 4 or later during the ‘Late Reproductive’ window. Data represent the mean and SEM. **C)** Box and whiskers plot of average population size (representing the number of F1 and F2 progeny) produced from a single founder *C. elegans*. Boxes indicate the 25-75th quartiles with median indicated. Error bars are the normalized range of the data and circles indicate outliers. The number (n) of experimental samples analyzed is indicated for each treatment. Stastistical significance is indicated when treat-ments differ. Treatments that share letters are not significantly different, whereas treatments with differing letters are stastically significant, post-hoc Dunn’s multiple comparison tests.

One explanation for the overall reduction in average brood size is the increased early mortality we observe in immunocompromised hosts. To address this possibility, we re-analyzed brood size only from hosts that survived past Day 3 and found that while this did slightly increase average brood size across all treatments, it was not significant between analyses (p=0.2356 two-tailed t-test) and exposure to *C. albicans* still significantly reduced average brood size (94±6) relative to uninfected (163±5) and heat-killed controls (141±9) (Figure S2, *p* < 0.0001 Kruskal-Wallis test). For hosts that survived past three days, we also detected a significant delay in reproductive timing upon exposure to *C. albicans*, with nearly 45% of all reproduction occurring in this late window (Figure 3C), similar to what we observe in healthy hosts (Table S1).

We predicted that lineage expansion in immunocompromised hosts exposed to *C. albicans* would be more dramatically impacted relative to wild-type nematodes exposed to *C. albicans*, given the reproductive delays coupled with the overall reduced brood sizes and increased mortality observed. Indeed, we determined that the median population size of *C. albicans* exposed nematodes was ∼11,000 compared to ∼17,000 and 19,500 in heat-killed and uninfected controls, respectively (Figure 3C). The overall reduction in population size resulting from *C. albicans* exposure is ∼43% in immunocompromised hosts and is significantly greater than the ∼32% reduction in population size in healthy hosts (Figure 4 and Table S1). Taken together, our results indicate that immunocompromised hosts are more susceptible to fungal infection than healthy wild-type hosts.

**Figure 4.**
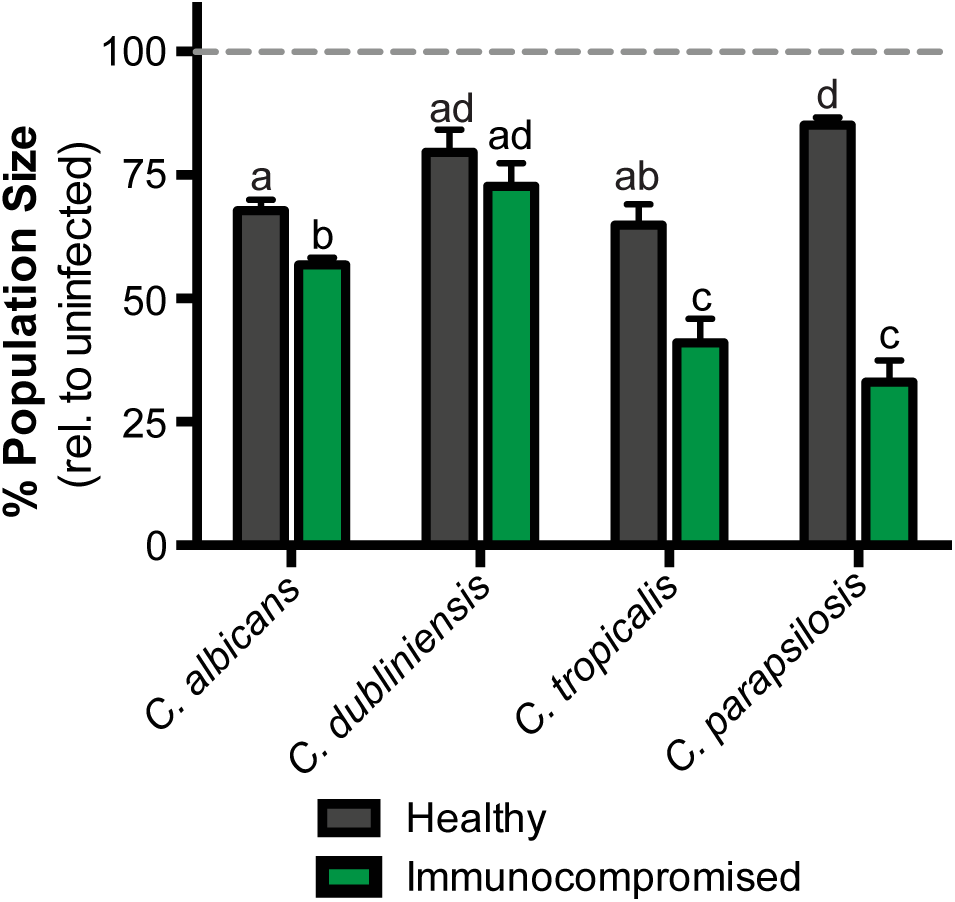
Virulence of *Candida* species depends on host immune function. Reductions in relative population size for healthy (N2, black bars) and immunocompromised (*sek-1*, green bars) hosts exposed to *C. albicans, C. dublineinsis, C. tropicalis*, and *C. parapsilosis*. Dashed line indicates the relative population size of uninfected hosts. Interactions between *Candida* species and host background was determined by two-way ANOVA (p < 0.001). Treatments that share letters are not significantly different, whereas treatments with differing letters are stastically significant, post-hoc Tukey’s multiple comparisons test. The number (n) of experimental samples analyzed is indicated for each treatment in Table S1.

### Host immune status reveals pathogenic potential in other Candida species

While *C. albicans* is the predominant fungal agent in invasive candidiasis, non-*albicans Candida* species (NACs) are estimated to account for 35-65% of candidaemia [41]. By evaluating NACs in both healthy and immunocompromised hosts, we can begin to assess their pathogenic potential, and thus extend the utility of *C. elegans* as a model host system for detecting virulence in *Candida* species. Here, we leveraged our simple, yet robust, lineage expansion assays to measure the impact of three additional *Candida* species: *C. dubliniensis, C. tropicalis* and *C. parapsilosis* on host fitness for healthy and immunocompromised *C. elegans*. All three non-*albicans Candida* species significantly reduced the population size of both healthy and immunocompromised hosts compared to uninfected controls (black asterisks, Figure S3 and Table S1), and sometimes were also significantly different than population sizes of hosts exposed to *C. albicans* (red asterisks, Figure S3).

We next wanted to determine if there was a significant interaction between *Candida* species and host immune function. To test this, we evaluated the population size of infected nematodes relative to uninfected for healthy (Figure 4; black bars) and immunocompromised (Figure 4; green bars) host backgrounds, since immunocompromised hosts have significantly smaller populations even when uninfected (Figure 2 and S4). There are significant differences among *Candida* species and nematode host backgrounds (p < 0.001, two-way ANOVA; p < 0.001 between pathogens, hosts, and their interaction). In healthy hosts, *C. albicans* and *C. tropicalis* are the most virulent, with 32-35% reductions in total population size, whereas *C. parapsilosis* and *C. dublineinsis* result in more modest population reductions (17-20%). However, in immunocompromised hosts, *C. parapsilosis* is one of the most virulent, with nearly a 67% reduction in population size. *C. albicans* and *C. tropicalis* are also highly virulent in immunocompromised hosts and all three of these species more severely impact immunocompromised hosts compared to healthy hosts. In contrast, *C. dublineinsis* is the least virulent, with no significant differences observed between the relative population sizes of healthy and immunocompromised hosts (Figure 4). Given the virulence phenotypes of these different yeasts, our results suggest that fungal virulence depends not only on the species of pathogen but also on the immune status of the host.

## Discussion

Here we utilized a *C. elegans* experimental host system to identify novel measures of host fitness associated with fungal infection. By tracking the number of progeny produced per day, we can quickly and quantitatively assess three aspects of host fitness: early mortality, total viable offspring produced and reproductive timing. We show that exposure to *Candia albicans* in healthy *C. elegans* hosts not only reduces survival (Figure 1), it modestly reduces total progeny produced, and dramatically delays reproductive timing (Figure 2). This reproductive delay has long-term consequences for lineage expansion, as single founder nematodes exposed to *C. albicans* reduces its population growth by ∼30% (Figure 2D). We found similar delays in reproductive timing in immunocompromised hosts exposed to *C. albicans*, although immunocompromised hosts are more susceptible to infection and have higher incidence of early mortality and smaller brood sizes compared to healthy hosts (Figure 3). Importantly, the delayed reproduction phenotype in infected *C. elegans* is easily ascertained, highly quantitative and reproducible and can be used as a screening method to detect relatively small differences in virulence across *Candida* strains or other fungal species. Furthermore, by utilizing different host contexts, this assay revealed the pathogenic potential of other important, yet understudied, *Candida* species (Figure 4), including *C. parapsilosis*, which has dramatically higher levels of virulence in immunocompromised hosts compared to healthy hosts.

The reductions in host fitness associated with *C. albicans* and other *Candida* species indicate that we have established a bona fide model of fungal infection. Importantly, these reductions in host fitness cannot be attributed to *Candida* species being a sub-optimal food source or that the hosts are starving because we deliver the pathogen with *E. coli*, the standard food source for *C. elegans*. Further, we do not detect any host larval development into dauer, an alternative lifestage in response to starvation and overpopulation [32]. While pathogen avoidance is a common defense strategy for *C. elegans* [42], we censored any worms that have crawled off the plate or have disappeared during the course of our experiment and removed them from our data analysis. Furthermore, the reductions in host fitness depend on ingesting live *C. albicans*, as heat-killed treatments do not cause significant reductions in total brood size, reproductive timing, or lineage expansion (Figure 2, ‘HK’). Microscopic analysis reveals that fungal cells inhabit the gut of *C. elegans* ([39] and data not shown) and we can extract viable host-associated fungal cells (data not shown). Taken together, these data support a model of fungal infection in the host *C. elegans*.

The host reproductive delay we observed in *C. elegans* upon fungal infection is a robust, highly quantitative measure of virulence that makes it amenable to screen a variety of host and pathogen genetic backgrounds that has been challenging in mammalian and insect models. *C. elegans* is easily maintained in the lab, has a short lifecycle that generates a large number of progeny, and has been a fundamental model genetic organism [32]. Previous work has shown that upon exposure to *C. albicans, C. elegans* have a transcriptional response consistent with that of infection [28]. In this work, we utilized a mutant *C. elegans* strain in which *SEK-1*, an important MAPK in innate immunity and whose homologs include the MKK3/6 and MKK4 family of mammalian MAPKKs, to determine the severity of fungal infection in immunocompromised hosts. We demonstrate that deficiencies in innate immune function result in hosts with high susceptibility to fungal infection and in the future can extend this analysis for many other host backgrounds that are mutant for immune function.

Previous studies using *C. elegans* as a host for fungal infection have described additional phenotypes beyond reduced survival, including a deformed anal region (DAR) [39,43]. It is possible that DAR may be contributing to the delayed reproductive phenotype we observe in our experiments, however, the frequency of DAR phenotypes was low and we did not observe any detectable differences between hosts exposed to *C. albicans* and hosts unexposed (data not shown), nor did we observe dramatic reductions in total number of viable offspring produced between these two groups of hosts (Figure 2A. While this deformity is a result of a local defense reaction of the worm due to extracellular signal-regulated kinase activation of the innate immunity MAKK cascade, it is a distinct marker from delayed reproduction for infection [44]. Wildtype reproductive timing utilizes the highly conserved DAF-2 DAF-16 insulin-like signaling pathway [23], but has also been implicated in innate immunity and bacterial infection [45] and is potentially disrupted in *C. elegans* hosts infected with *Candida*.

Impacts on host reproduction have often been overlooked in studies regarding *Candida* virulence, yet host reproduction is an important evolutionary measure of an organism’s fitness. Not only can we utilize these fecundity assays to assess the roles of host immune function on fungal pathogenesis, we also readily screen diverse pathogen strain backgrounds and species. A large-scale, international 10-year study identified 31 species of *Candida* associated with clinical samples. While *C. albicans* is still the most prevalent, its isolation from clinical isolates is decreasing with corresponding increases isolation of *C. glabrata, C. tropicalis* and *C. parapsilosis* [46]. Despite its close evolutionary relationship with *C. albicans, C. dubliniensis* does not seem to be highly pathogenic [47]. However, other non-*albicans Candida* species have been increasingly implicated in fungal infections of humans [48]. For example, *C. tropicalis* and *C. parapsilosis* are common fungal species isolated in patients with candidemia (7 - 48% and 11 - 27% respectively and depending on geographic region), candiduria (8 – 44% and 0.5 – 11% respectively, and oral candidosis (5 – 13% and 7 – 15%, respectively) ([48] and references therein). Despite the increasing incidence of non-*albicans Candida* infections [46], experimental studies using these pathogens remains limited. Here, we analyzed the virulence phenotypes of three non-albicans *Candida* species, *C. dubliniensis, C. tropicalis* and *C. parapsilosis* in both healthy and immunocompromised host backgrounds. In healthy *C. elegans* hosts, *C. parapsilosis* is the least virulent compared to *C. albicans* and *C. tropicalis* and *C. dublineinsis* has intermediate level of virulence (Figure 4). For all species except for *C. dubliniensis*, virulence is more severe in immunocompromised hosts and is particularly striking in *C. parapsilosis*, which is the most virulent in hosts with compromised immune function. Therefore, the *C. elegans* fecundity assays we developed in this study can rapidly reveal pathogenic potential by assessing virulence in multiple host genetic backgrounds and across pathogens in a highly quantitative and robust manner.

## Supporting information

