## Supplementary material for "A novel virulence phenotype rapidly assesses *Candida* fungal pathogenesis in healthy and immunocompromised *Caenorhabditis elegans* hosts"

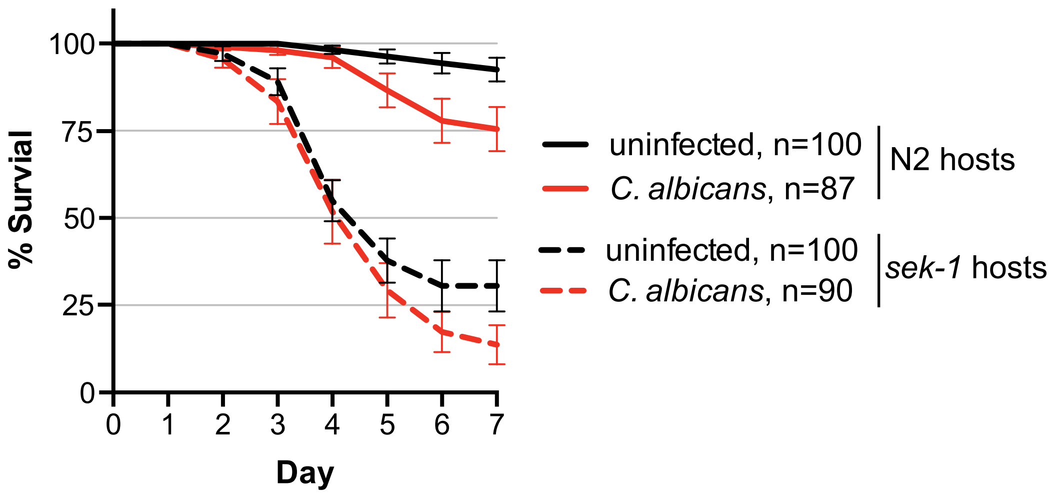


**Figure S1: Immunocompromised hosts survive less well than healthy hosts.** Mean survival of healthy N2 (solid lines) and immunocompromised *sek-1* (dashed lines) *C. elegans* populations that are either uninfected (black) or when infected with *C. albicans* SC5314 (red). Mean data values are plotted with SEM error bars, number of worms analyzed (n) for each treatment is indicated in the legend.
