## Supplementary material for "A novel virulence phenotype rapidly assesses *Candida* fungal pathogenesis in healthy and immunocompromised *Caenorhabditis elegans* hosts"

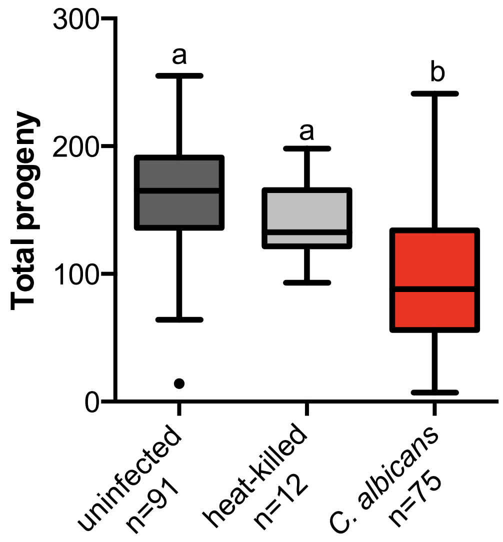


**Figure S2: Reduced brood size in *C. albicans*-infected immunocompromised hosts is not due to early death.** Box and whiskers plot of brood sizes for immunocompromised *C. elegans* (*sek-1*) that are still alive on or by Day4 that have been exposed to *E. coli* food source alone (uninfected, black) or live *C. albicans* (red). Boxes indicate the 25-75th quartiles with median indicated. Error bars are the normalized range of the data and circles indicate outliers. The number (n) of experimental samples analyzed is indicated for each treatment. Treatments that share letters are not significantly different, whereas treatments with differing letters are stastically significant, post hoc Dunn’s multiple comparison test.
