## Supplementary material for "A novel virulence phenotype rapidly assesses *Candida* fungal pathogenesis in healthy and immunocompromised *Caenorhabditis elegans* hosts"

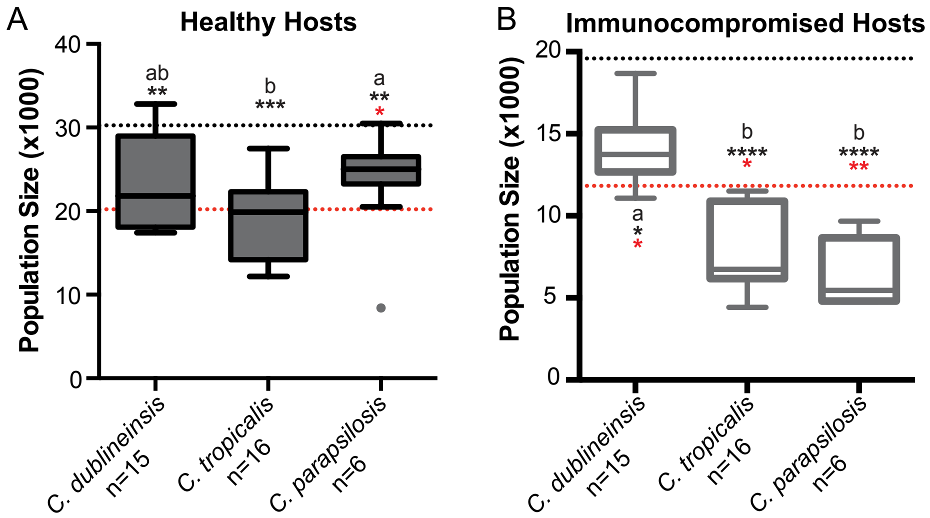


**Figure S3: Non-*albicans Candida* species differentially impact host fitness depending on host immune status.** Box and whiskers plot of average population size from lineage expansion assays (representing the number of F1 and F2 progeny) produced from a single founder *C. elegans* for A) healthy hosts (N2) or B) immunocompromised hosts (*sek-1*) . exposed to *C. dublineinsis, C. tropicalis and C. parapsilosis*. Boxes indicate the 25-75th quartiles with median indicated. Error bars are the normalized range of the data and circles indicate outliers. The number (n) of experimental samples analyzed is indicated for each treatment. Treatments that share letters are not significantly different, whereas treatments with differing letters are stastically significant, post-hoc Dunn's multiple comparisons test. Mean brood size for uninfected (black line) and *C. albicans* exposed (red line) hosts is indicated for reference. Astersisks indicate significant difference from uninfected (black) or live *C. albicans* (red) for each non-*albicans Candida* species (post-hoc Dunn's multiple comparison tests).
