## Supplementary material for "A novel virulence phenotype rapidly assesses *Candida* fungal pathogenesis in healthy and immunocompromised *Caenorhabditis elegans* hosts"

| | | uninfected | heat-killed *C. albicans* | Live *C. albicans* | *C. dubliniensis* | *C. tropicalis* | *C. parapsilosis* |
|---|---|---|---|---|---|---|---|
| Brood Size | Healthy (N2) | 285±4 (n=99) | 281±6 (n=26) | 253±7 (n=95) | 276±17 (n=19) | 260±5 (n=9) | 224±9 (n=18) |
| | Immunocompromised (sek-1) | 151±6 (n=102) | 127±13 (n=14) | 78±6 (n=93) | 94±17 (n=16) | 105±16 (n=15) | 38±13 (n=14) |
| | p-value | **** | **** | **** | **** | **** | **** |
| % Late Reproduction | Healthy (N2) | 19±1.2 (n=99) | 14±2.7 (n=26) | 44±1.7 (n=93) | 21±3.7 (n=19) | 32±7.2 (n=9) | 72±13 (n=8) |
| | Immunocompromised (sek-1) | 26±1.5 (n=90) | 24±2.8 (n=12) | 46±2.2 (n=75) | 40±4.7 (n=13) | 30±3.4 (n=6) | 30±3.4 (n=6) |
| | p-value | *** | 0.042 | 0.355 | ** | 0.758 | **** |
| Lineage Expansion | Healthy (N2) | 29481±756 (n=31) | 30319±1181 (n=12) | 19964±636 (n=16) | 23492±1358 (n=15) | 19115±1271 (n=16) | 24437±799 (n=9) |
| | Immunocompromised (sek-1) | 19417±549 (n=23) | 17342±599 (n=11) | 11043±277 (n=35) | 14107±909 (n=7) | 7917±950 (n=8) | 6417±852 (n=6) |
| | p-value | **** | **** | **** | **** | **** | **** |

**Table S1:** Brood size, fraction of late reproduction, and lineage expansion population size of healthy (N2) and immunocompromised (sek-1) hosts when treated with food alone (uninfected), heat-killed (MH88) or live *Candida* species. Data is the average +/- SEM, and the total number of worms analyzed (n) is provided. P-values are given for two-tailed t-tests comparing healthy and immunocompromised individuals for each treatment.
